# Charge-Driven Fibril Recognition and Covalent Disruption of A*β*_42_ by Paddlewheel Diruthenium Complexes

**DOI:** 10.64898/2026.06.26.734728

**Authors:** Alejandro Feito, Andrés R. Tejedor, Alberto Ocana, Aarón Terán, Antonello Merlino, Daniela Marasco, Santiago Herrero, Jorge R. Espinosa

## Abstract

The inhibition of A*β*_42_ (*β*-amyloid) fibril formation is a key therapeutic strategy in Alzheimer’s disease research. Paddlewheel diruthenium complexes have shown promising activity against A*β*_42_ aggregation and preformed fibril disaggregation, yet their molecular mode of action remains poorly understood. In this work, we perform atomistic simulations to explore how charge modulation influences the interactions of three analogous paddlewheel diruthenium complexes, the parent neutral complex [Ru_2_Cl(D-*p*-FPhF)(O_2_CCH_3_)_3_], and its anionic [Ru_2_Cl_2_(D-*p*-FPhF)(O_2_CCH_3_)_3_]^*−*^ and cationic [Ru_2_(D-*p*-FPhF)(O_2_CCH_3_)_3_]^+^ counterparts (D-*p*-FPhF^*−*^ is the *N,N’* -bis(4-fluorophenyl)formamidinato ligand) with A*β*_42_. Our results indicate that electrostatic tuning governs binding affinity and the extent of interaction across the A*β*_42_ fibril surface. As the complexes’ charge changes from -1 to +1, the interaction pattern shifts from localized contacts to widespread, multi-site engagement encompassing key charged, aromatic, and hydrophobic regions of A*β*_42_. This enhanced binding correlates with longer-lived, thermodynamically stable interactions at the fibril interface, which effectively lower the free energy penalty for fibril disassembly. Overall, our findings propose a mechanism in which charge-dependent activation through ligand exchange enhances fibril recognition and promotes disruptive binding modes, demonstrating the potential of charge-tunable diruthenium complexes as therapeutic modulators of A*β*_42_ fibril stability.

## INTRODUCTION

Alzheimer’s disease (AD) is the most prevalent neurodegenerative disorder worldwide and the leading cause of dementia, currently affecting more than 55 million individuals globally and is expected to triple, with projections exceeding 150 million cases by 2050.^1,2^ Despite decades of intensive research, available treatments remain purely symptomatic and are unable to halt or reverse disease progression.^3^

From a neuropathological perspective, AD is defined by two hallmark histological features: intracellular neurofibrillary tangles composed of hyperphosphorylated tau protein and extracellular senile plaques primarily formed by the *β*-amyloid (A*β*) peptide.^1^ According to the amyloid cascade hypothesis,^4^ the abnormal production, aggregation, and deposition of A*β* constitute the primary pathogenic event triggering neurodegeneration. A*β* is generated through sequential cleavage of the amyloid precursor protein (APP) by *β*- and *γ*-secretase, producing peptide isoforms ranging from 37 to 43 amino acids, among which A*β*_1–40_ and A*β*_1–42_ are the most abundant.^2^

Among these isoforms, A*β*_1–42_ is particularly relevant from a pathogenic standpoint, as its critical aggregation concentration is approximately fivefold lower than that of A*β*_1–40_, rendering it significantly more prone to the formation of oligomers, protofibrils, and mature amyloid fibrils.^5^ Although early studies identified insoluble mature fibrils as the principal neurotoxic species (fibril hypothesis), growing evidence indicates that soluble oligomers could represent the most cytotoxic intermediates, capable of disrupting synaptic signaling and inducing neuronal death independently of fibrillar plaque formation.^1,3^ Although inhibition of fibrillar aggregation remains a major therapeutic goal, as fibrils can act as seeds that accelerate polymerization through nucleation-dependent mechanisms,^5^ the ability to clear insoluble A*β* seems to be a more important factor than preventing the A*β* aggregation.^6–8^

The repeated failure of clinical trials based on anti-A*β* monoclonal antibodies has stimulated the exploration of alternative therapeutic strategies.^1^ In this context, metallodrugs have emerged as promising candidates owing to the ease with which their coordination geometry, oxidation state, lipophilicity, and ligand-exchange kinetics can be tuned. Indeed, these parameters directly govern how a metal complex interacts with biomolecular targets such as A*β*.^9,10^ Among metal-based systems displaying antiamyloidogenic activity, paddlewheel diruthenium complexes Ru_2_^5+^ stand out for their metal–metal multiple bond, aqueous solubility and stability, distinct coordination environments (four equatorial and two axial sites, see Figure 1) and the easy modulation of the steric and electronic properties.^11–13^ These features have no equivalent in organic molecules or in mononuclear metal complexes, and it is precisely what gives the Ru_2_^5+^ core its distinctive reactivity: the axial positions provide a kinetically accessible entry point for a first encounter with a biomolecule,^14^ while the equatorial carboxylates offer a separate, more controlled site for subsequent ligand exchange.^15^ X-ray crystallographic studies of diruthenium adducts with model proteins, such as hen egg-white lysozyme (HEWL) and bovine pancreatic ribonuclease A (RNase A), have provided structural evidence for this two-step sequence at atomic resolution: the labile axial ligand is displaced first, typically by His, Lys, or Arg side chains, after which an equatorial carboxylate is lost in favor of an Asp or Glu residue — with the Ru–Ru bond and paddlewheel topology preserved throughout.^16–19^

**FIG. 1.**
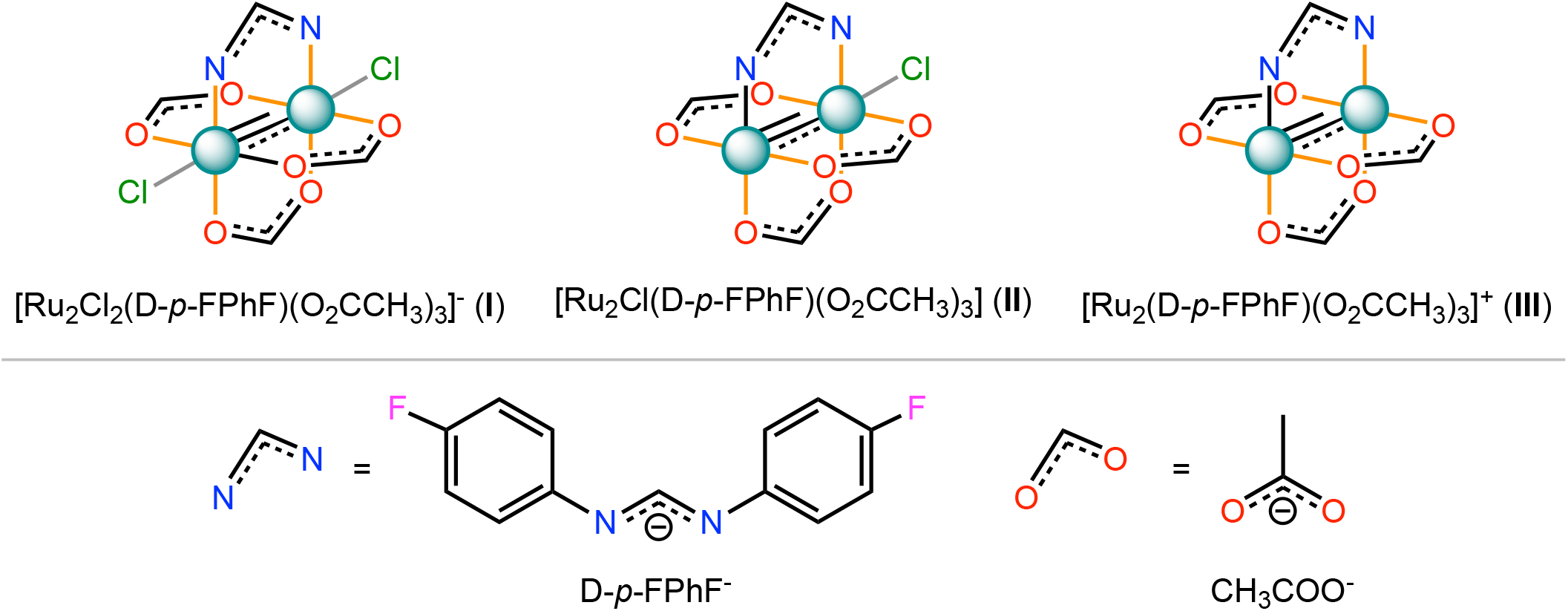
Structure of the paddlewheel diruthenium complexes I, II and III studied in this work. Grey and orange bonds indicate the axial and equatorial ligands, respectively, in these species.

This reaction sequence has been established for globular proteins, but whether the same axial-then-equatorial logic governs recognition of the structurally distinct surface of an amyloid fibril remains an open question. The [Ru_2_Cl(D-*p*-FPhF)(O_2_CCH_3_)_3_] compound (D-*p*-FPhF^*−*^ = *N,N’* -bis(4-fluorophenyl)formamidinato) offers a system to address this question, since it has already been shown to act as a potent inhibitor of A*β*_1–42_ fibril formation, redirecting the peptide toward non-toxic oligomeric states, disaggregating preformed fibers, and fully suppressing peptide-induced cytotoxicity.^11^ This activity was attributed to a charge- and steric-dependent mode of binding of the metal complex to the peptide chain,^11,12,17^ but the experimental data could not resolve the atomistic sequence of events, nor how that charge state governs the transition from a localized initial contact to multi-site binding of the fibril surface that is ultimately responsible for its disruption.

Elucidating the atomistic determinants of metallo-drug–biomolecule recognition at the molecular level remains a formidable challenge for purely experimental approaches, owing to the transient nature of the binding events and the heterogeneity of the binding-competent conformations^20^. Molecular dynamics (MD) simulations have emerged as a powerful complementary tool, providing residue-level resolution of contact patterns, binding site dynamics, and free energy landscapes that are inaccessible to bulk spectroscopic or biochemical assays.^21–26^ In the context of inorganic and metallodrug–peptide interactions, MD have been successfully employed to probe the binding modes of platinum complexes to DNA and to model their interactions with proteins,^27^ to map the interaction surfaces of ruthenium-based anticancer agents with serum albumin and nucleobases,^28,29^ and more recently to resolve the conformational consequences of metal coordination on intrinsically disordered peptides relevant to neurodegeneration.^30**?**^ Despite these advances, the interplay between charge state, ligand exchange, and fibril-surface recognition for paddlewheel diruthenium complexes has not been characterized at atomic level.

Here, we investigate the interaction between A*β*_42_ fibrils^31,32^ and three analogous paddlewheel diruthenium complexes, the parent neutral complex [Ru_2_Cl(D-*p*-FPhF)(O_2_CCH_3_)_3_],^11,13^ and its anionic [Ru_2_Cl_2_(D-*p*-FPhF)(O_2_CCH_3_)_3_]^*−*^ and cationic [Ru_2_(D-*p*-FPhF)(O_2_CCH_3_)_3_]^+^ counterparts (Figure 1), to discern how the charge of these species can affect the metal-amyloid fibril recognition. Thus, we use extensive all-atom MD simulations, free energy calculations, and residue-level contact analysis to assess how axial substitution modulates the interaction with the fibrillar assembly^12^. Our analysis focuses on residue-specific contact maps, free energy profile reconstruction of fibril dissociation, and the mechanistic consequences of complex–fibril adduct formation. By systematically varying the axial coordination environment, we demonstrate that the difference in chloride ligand content acts as a charge-activation mechanism that enhances fibril recognition, while subsequent covalent coordination to an Asp23 residue acts as a structural locking step that locally disrupts the cross-*β* architecture. Our results provide a molecular-level rationale for the experimentally observed anti-amyloidogenic activity of diruthenium compounds^11,12^.

## METHODOLOGY

We performed all-atom MD simulations employing the a99SB-*disp* force field together with the TIP4P-*disp* water model^33–36^. This combination has been shown to provide a balanced description of both folded and intrinsically disordered proteins^33,37^, which is particularly important for accurately capturing conformational fluctuations and ligand recognition events in biomolecular systems. The parameters for the diruthenium complexes were obtained from density functional theory (DFT) calculations performed with Gaussian 09^38^ using the B3LYP-D3/def2-TZVP functional/basis set combination. The derived bonded and electrostatic parameters were incorporated into the a99SB-*disp* framework, ensuring full consistency with the all-atom representation of the protein interactions^39^.

All simulations were performed using GROMACS (version 2023)^40^ at physiological conditions (150 mM NaCl), at 300 K and at 1 bar pressure. Our systems were solvated in cubic boxes of 7.5 × 7.5 × 7.5 nm, where water molecules were replaced with Na^+^ and Cl^*−*^ ions to neutralize the systems and reach the target salt concentration. Energy minimization was conducted using a force convergence criterion of 1000 kJ mol^*−*1^ nm^*−*1^. All bonds involving hydrogen atoms were constrained using the LINCS algorithm,^41^ allowing a 2 fs integration time step. Periodic boundary conditions were applied in all directions, and long-range electrostatics were treated with the Particle-Mesh Ewald (PME) method^42^ using a real-space cutoff of 0.9 nm. Temperature was controlled with the velocity-rescale thermostat (*τ*_*T*_ = 1.0 ps), and pressure was maintained using the Parrinello–Rahman barostat (*τ*_*P*_ = 1.0 ps). Each system was simulated for a total accumulated time of 500 ns.

For potential of mean force (PMF) calculations, systems were solvated in boxes of 7 × 7 × 10 nm at 300 K and 150 mM NaCl. The protein–ligand center-of-mass (COM) distance was biased using a harmonic umbrella potential with a force constant of 25000 kJ mol^*−*1^ nm^*−*2^. Approximately 40 umbrella windows were generated at 0.025 nm intervals along the reaction coordinate. During production runs, positional restraints of 1000 kJ mol^*−*1^ nm^*−*2^ were applied (perpendicular to the pulling axis) to heavy atoms of selected protein chains to prevent over-all rotation. Each umbrella window contributed to an accumulated simulation time totaling 400 ns. Free energy profiles were reconstructed using the Weighted Histogram Analysis Method (WHAM)^43^ as implemented in GROMACS. The first 2000 ps of each trajectory were discarded prior to analysis to ensure equilibration. Statistical uncertainties were estimated via block averaging, dividing each trajectory into four independent segments and computing PMF profiles separately for each block.

Furthermore, intermolecular contact maps within protein fibrils were computed from the all-atom trajectories using a distance-based criterion. Since relative contact frequencies are generally robust to reasonable variations in the cut-off distance, we adopted a sequence-dependent threshold based on excluded volume considerations. Specifically, contacts were defined when interatomic distances were below 1.2*σ*_*ij*_, where *σ*_*ij*_ corresponds to the average excluded volume parameter of amino acids *i* and *j*.^24,44,45^ For the heavy atoms of the diruthenium complexes, a *σ* value of 5 Å was assigned. Given that the minimum of the Lennard–Jones potential occurs at approximately 2^1*/*6^*σ*_*ij*_ (*∼* 1.122*σ*_*ij*_), the chosen 1.2*σ*_*ij*_ threshold ensures that only energetically meaningful contacts are considered^25,46^. Contact lifetime is computed directly from the trajectories as the time-averaged probability of being in the bound state. Specifically, for each ruthenium atom a contact function *h*(*t*) is defined (1 if within the cut-off distance to the protein and 0 otherwise), and then obtained as 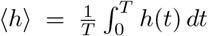, the fraction of simulation snapshots in which a contact is present.

## RESULTS AND DISCUSSION

### Structural characterization of the A*β*_42_ fibril and key residues governing fibril stabilization

The structural basis of A*β*_42_ fibril formation was analyzed using the PDB structure 2MXU,^31^ which comprises residues 11–42 of the A*β*_1*−*42_ peptide. This region encompasses the core amyloid-forming segment identified in previous experimental studies as essential for fibril nucleation and propagation (region 21–42).^11^ Within the A*β*_11*−*42_ fragment, the charged residues Glu11, Glu22, Asp23, Lys16 and Lys28, together with the aromatic residues Phe18 and Phe19 and the histidine residues His13 and His14 are spatially clustered at positions that are likely to contribute to intermolecular contacts stabilizing the fibril architecture (Figure 2A). These residues adopt well-defined spatial positions within the *β*-hairpin conformation characteristic of the fibrillar state (Figure 2B), whose repetitive stacking along the fibril axis gives rise to a highly ordered cross-*β*-sheet architecture (Figure 2C).

**FIG. 2.**
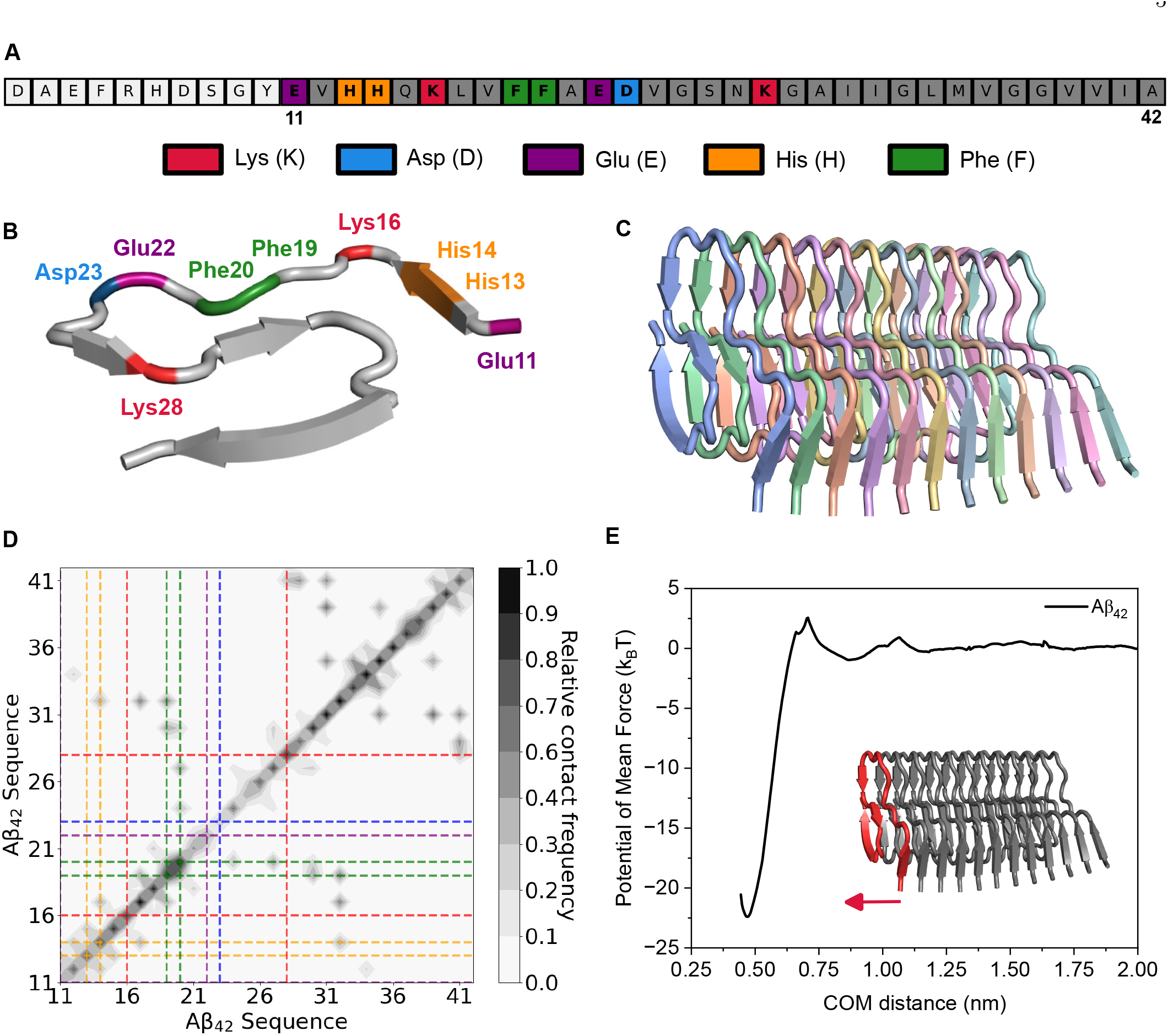
Structural and energetic characterization of the A*β*_42_ segment studied in this work (LARK segment). (A) Primary sequence of A*β*_1*−*42_ highlighting charged (Lys, Asp, Glu, His) and aromatic (Phe) residues within the LARK segment. (B) Representative conformation of the isolated LARK segment showing key charged and aromatic residues that participate in intermolecular contacts. (C) Parallel in-register *β*-sheet stacking arrangement of the hexameric LARK assembly. (D) Relative intermolecular residue–residue contact frequency map for A*β*_42_ within the fibril, with dashed lines marking the boundaries of the LARK segment. (E) Potential of mean force (PMF) as a function of the center-of-mass (COM) distance between stacked LARK segments, with a representative bound configuration at the free-energy minimum shown in the inset and the pulling segment in red.

The contact frequency map shown in Figure 2D from an unbiased MD simulation delineates the network of inter-residue interactions that stabilize the A*β*_42_ fibril. Two major classes of contacts are evident: (i) diagonal contacts, corresponding to in-register parallel interactions between identical residues on adjacent chains, and (ii) off-diagonal contacts, reflecting interactions between residues that are distant in sequence but proximal in three-dimensional space.

The strongest diagonal contacts are predominantly localized within the hydrophobic C-terminal region. Residues Leu34, Ile32, Met35, Ile31, and Leu17 exhibit the highest contact frequencies, indicating persistent pairing with their counterparts on neighboring chains. This pattern is characteristic of a parallel, in-register *β*-sheet architecture, in which hydrophobic side chains stack along the fibril axis. Notably, the clustering of contacts within the Ile31-Met35 segment highlights its central structural role. This region forms the C-terminal *β*-strand of the hairpin and constitutes a buried hydrophobic nucleus within the fibril core.^32^ In-register stacking of residues such as Ile32, Leu34, and Met35 across adjacent chains has been directly observed by intermolecular solid-state NMR.^32^ The prominence of Leu17 among the strongest diagonal contacts further supports the contribution of the N-terminal *β*-strand to inter-chain registry. This is consistent with mutagenesis studies showing that the central hydrophobic cluster (residues 17-21) is essential for A*β*_42_ aggregation and deposition.^47^

Off-diagonal contacts provide complementary insight into the tertiary packing within and between *β*-hairpin units. The most frequent interactions included Ile31-Val39, Phe19-Ile32, Ile31-Met35, Leu17-Ile32, and Ile31-Ile41. Contacts such as Ile31-Val39 and Ile31-Ile41 reported on cross-strand interactions across the hairpin turn, indicate tight apposition of the *β*-strands and effi-cient hydrophobic core packing. These observations are consistent with intramolecular contacts involving Ile31, Val36, Val39, and Ile41 identified by solid-state NMR in the fibril structure.^32^

To quantify the thermodynamic stability of the fibril and the energetic cost of removing a single chain from the assembled structure, we computed the potential of mean force (PMF) as a function of the center-of-mass (COM) distance between one terminal chain and the rest of the fibril (Figure 2E). The depth of the free energy well, which corresponds to the dissociation free energy of a single chain from the fibril, is approximately 22 *k*_B_*T*. This value is consistent with the strong thermodynamic stability typically associated with amyloid fibrils and is in quantitative agreement with estimates reported in previous computational studies of related peptide cross-*β*-sheet structures and RNA G-quadruplexes.^23,48–50^ Notably, the PMF does not show strong secondary minima or intermediate metastable states along the dissociation pathway, possibly in part due to the applied positional restraints^36^, suggesting that chain removal may proceed as a cooperative, single-step process without stable partially-dissociated intermediates. This cooperativity is mechanistically linked to the network of contacts that must be broken concurrently for dissociation to occur, which is a hallmark of the cross-*β* architecture.

In summary, our structural and energetic analyses provide a coherent molecular picture of A*β*_42_ fibril stabilization. The contact frequency map reveals a clear partitioning between diagonal, in-register intermolecular interactions that propagate the parallel cross-*β* architecture along the fibril axis, and off-diagonal contacts that reinforce the intramolecular *β*-hairpin fold. The hydrophobic C-terminal segment, particularly residues within the Ile31–Met35 region, emerges as the principal structural nucleus, supported by contributions from the central hydrophobic cluster around Leu17–Phe19. In contrast, charged residues such as Glu22 and Asp23 display minimal contact participation, consistent with a more solvent-exposed and structurally permissive role. Thermodynamically, the PMF profile demonstrates a substantial dissociation free energy barrier. Together, these results underscore the dominant role of hydrophobic, hydrogen-bonding, and aromatic packing in defining both the structural integrity and the cooperative stability of the A*β*_42_ cross-*β*-sheet assembly.

### Interaction of diruthenium complexes with A*β*_42_: effect of electrostatic interactions on the residue-specific binding pattern

Having established the key structural and energetic determinants of A*β*_42_ fibril stability, we next examined the binding of three diruthenium complexes with different charge to the fibril surface. The three complexes, [Ru_2_Cl_2_(D-*p*-FPhF)(O_2_CCH_3_)_3_]^*−*^ (Complex I, anionic), [Ru_2_Cl(D-*p*-FPhF)(O_2_CCH_3_)_3_] (Complex II, neutral), and [Ru_2_(D-*p*-FPhF)(O_2_CCH_3_)_3_]^+^ (Complex III, cationic), differ in their axial ligand environment, which progressively modulates the net charge of the complexes. All simulations were performed using 12 A*β*42 chains and 12 diruthenium complexes, maintaining a 1:1 stoichiometry between peptide and complex, consistent with the setup employed by La Manna *et al*.,^11^. This design is motivated by the known lability of axial chloride ligands in this class of diruthenium systems, which under physiological conditions can undergo dissociation and subsequent solvent coordination.^51^ Consequently, exploring different coordination complexes allows us to assess which one maximizes interactions with the fibrillar interface. The residue-specific contact frequency maps between each complex and the A*β*_42_ sequence (residues 11– 42) are shown in Figure 3A–C, where the colored dashed lines indicate the positions of key residues identified in the fibril analysis: charged residues Lys16 and Lys28 (red), Asp23 (blue), Glu11 and Glu22 (violet), aromatic residues Phe19 and Phe20 (green), and histidine residues His13 and His14 (orange).

**FIG. 3.**
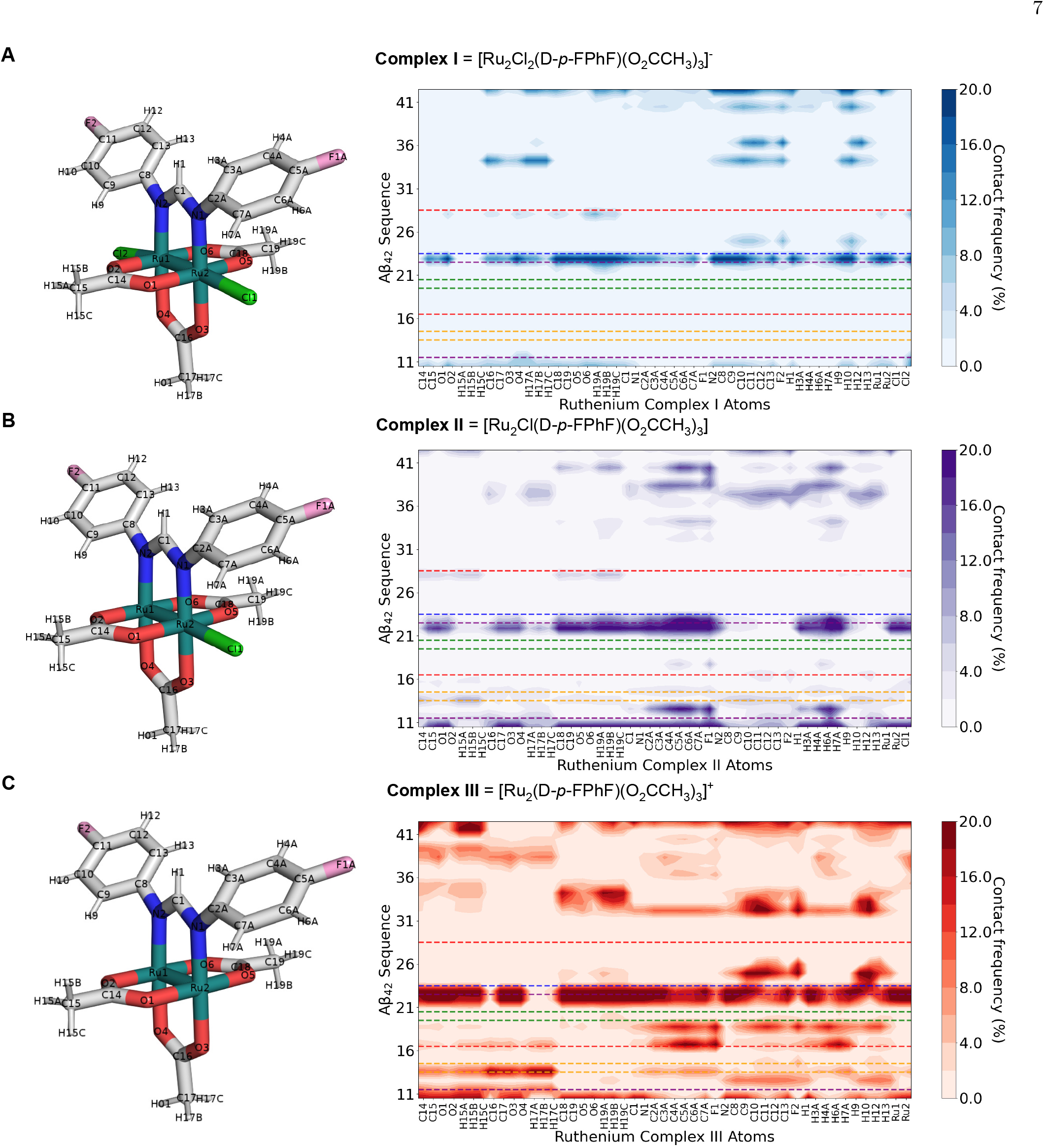
Interaction analysis between A*β*_42_ and three diruthenium complexes. (A–C, left) Structures of the complexes [Ru_2_Cl_2_(D-*p*-FPhF)_2_(O_2_CCH_3_)_3_]^*−*^ (Complex I), [Ru_2_Cl(D-*p*-FPhF)(O_2_CCH_3_)_3_] (Complex II), and [Ru_2_(D-*p*-FPhF)(O_2_CCH_3_)_3_]^+^ (Complex III). (A–C, right) Residue–atom contact frequency maps (%) between A*β*_42_ and each diruthenium complex, showing the relative interaction probability across the peptide sequence. Horizontal dashed lines highlight key sequence regions of interest. Color scales indicate the percentage of contacts formed during the simulation of the fibril in presence of the complexes.

Complex I, bearing two axial chloride ligands and carrying a negative charge, displays a sparse and localized contact pattern (Figure 3A). Detectable interactions are predominantly confined to the central region of the peptide, with modest contact frequencies concentrated around residue Asp23. The N-terminal region (residues 11-16) and the hydrophobic C-terminal segment (residues 30-42) are largely devoid of significant contacts. This restricted binding footprint is consistent with the absence of strong electrostatic driving forces in the anionic complex, which limits interactions to those of hydrophobic or van der Waals character and reduces the accessible binding surface on the fibril. This sparse, non-covalent mode of engagement parallels the behavior reported crystallographically for anionic diruthenium species, such as [Ru_2_(CO_3_)_4_]^3*−*^ and [Ru_2_(L-L)(CO_3_)_3_]^2*−*^ (where L-L is a formamidinate ligand), which remain electrostatically associated with the surface of HEWL without displacing equatorial ligands, as long as the complex retains its negative charge in solution.^18,19^ Loss or substitution of a carbonate ligand in those systems reduces the net charge and switches the binding mode to covalent coordination, indicating that the surface-confined footprint of Complex I reflects a general property of anionic Ru_2_^5+^ species rather than a feature specific to the amyloid fibril surface.

Complex II, bearing one axially coordinated chloride ligand, acquires neutral charge and exhibits a notably more extensive contact pattern (Figure 3B). While the region around residues Glu22-Asp23 retains significant contact frequency, interactions now extend more broadly along the peptide sequence, with emerging contacts close to Glu11. The appearance of contacts with His residues is particularly noteworthy, as the imidazole side chains of His13 and His14 are known metal-coordinating groups that can engage in direct interactions with diruthenium centers.^11^ The emergence of these contacts is consistent with the role of histidine as an axial anchoring residue for the Ru_2_ core, observed crystallographically in the adduct formed between the neutral compound [Ru_2_(D-*p*-FPhF)(O_2_CCH_3_)_2_(O_2_CO)] and RNase A.^52^ The partial extension of contacts toward the N-terminal hydrophobic region also becomes apparent, suggesting that the neutral charge facilitates access to binding sites that were sterically or energetically inaccessible to the anionic complex.

Complex III, without axial chloride ligands, carries a positive charge and presents a dramatically broadened and intensified contact frequency map (Figure 3C). High-frequency contacts now involve nearly the entire A*β*_42_ sequence from residue 11 to 42, with the most intense interactions distributed across multiple regions simultaneously. The negatively charged residues Glu11, Glu22, and Asp23 all show substantial contact frequencies, as do the aromatic residues Phe19 and Phe20 and the histidine pair His13-His14, consistent with crystallographic evidence for His-axial coordination by cationic diruthenium fragments, and computational ev-idence for chloride loss generating the reactive cationic species.^16^ Strikingly, the C-terminal hydrophobic core (residues 30-42), which harbors the principal structural nucleus of the fibril as characterized above, now displays marked contacts with the complex. This broad engagement with the fibril surface suggests that the cationic complex is capable of interacting simultaneously with multiple residue classes —charged, aromatic, and hydrophobic—thereby disrupting the cooperative network of contacts that sustains the cross-*β*-sheet architecture. In the crystal structure of the adduct formed between [Ru_2_Cl(D-*p*-CNPhF)(O_2_CCH_3_)_3_] (D-*p*-CNPhF^*−*^ = *N,N’* -bis(4-cyanophenyl)formamidinate) and RNase A, the diruthenium core binds the His105 side chain at the axial site.^16^ Quantum chemical calculations support the idea that the cationic complex formed by replacing the chloride axial ligand by a water molecule is the species responsible of interacting with this residue.^16^ The prominence of Asp23 as the dominant equatorial anchor for Complex III is consistent with the recurrence of Asp residues (*e*.*g*. Asp101 or Asp119) as the principal equatorial binding sites identified across different cationic diruthenium–HEWL adducts previously reported,^16–19^ suggesting that Asp carboxylates represent a conserved equatorial recognition motif for cationic Ru_2_ fragments independently of the host biomolecule, even when that biomolecule lacks the folded tertiary scaffold of HEWL or RNase A.

Taken together, the progression from Complex I to Complex III reveals a clear and systematic dependence of the binding footprint on the overall charge of the diruthenium complex. As the complex global charge changes from negative to positive, the contact pattern evolves from a localized interaction, primarily centered on the Glu22-Asp23 region, to a broad multi-site engagement that spans charged, aromatic, and hydrophobic residues distributed across the entire A*β*_42_ sequence. This charge-driven expansion of the binding interface indicates that electrostatic attraction between the cationic complex and the negatively charged and polar residues of the fibril surface is the primary determinant of binding affinity and selectivity. Importantly, Complex III reproduces the residue-level interaction pattern most consistent with the experimentally identified fibril contact map reported previously,^11^ including significant interactions with residues involved in both the central hydrophobic cluster and the C-terminal *β*-strand that stabilizes the cross-*β* architecture. This agreement suggests that the cationic complex provides the most realistic description of the fibril-binding form of the complex.

Our results further support a mechanistic scenario in which axial chloride dissociation generates a coordinatively unsaturated and highly reactive species capable of engaging the fibril surface. In this state, the exposed coordination sites can directly interact with key fibril residues, thereby competing with native inter-chain contacts. In particular, the ability of Complex III to target the hydrophobic C-terminal nucleus—the region most critical for fibril stability—suggests a potential disruption mechanism based on direct competition with the intermolecular interactions that propagate the parallel *β*-sheet along the fibril axis.

To further quantify the residue-specific binding preferences revealed by the contact frequency maps, we ranked the top ten A*β*_42_ residues by their relative contact frequency with each complex (Figure 4A). For Complex I, the dominant interaction involves Asp23, which accounts for the maximum normalized contact frequency (1.000), followed by Ala42 and Glu11 at considerably lower values. The predominance of a single negatively charged residue at the top of the ranking is consistent with the sparse, localized binding footprint described above and underscores the limited chemical diversity of the binding interface accessible to the anionic complex. The ranking for Complex II reveals a qualitative shift: Glu11 now emerges as the top-ranked residue, and the list includes His13 and His14, both of which carry metal-coordinating imidazole side chains. The emergence of histidine residues in the ranking supports the notion that the neutral Complex II enables direct coordination or close approach to nitrogen-donor sites that are not significantly engaged by the anionic species. The broader distribution of contact frequencies across the top ten residues further corroborates the expanded binding footprint inferred from the contact maps. For Complex III, the ranking reflects the most chemically diverse and spatially distributed binding profile among the three complexes. Asp23 again ranks first (1.000), but is now accompanied by Glu22, Glu11, and, strikingly, aromatic and hydrophobic residues including Phe19 and Ile32, as well as His14. The simultaneous presence of charged, aromatic, and aliphatic residues among the top-ranked contacts indicates that Complex III engages all major classes of fibril surface residues. Notably, the residues appearing in the top ten for Complex III—including Glu11, Glu22, Asp23, His13, His14, Phe19, and hydrophobic core residues—closely match the interaction pattern identified experimentally by La Manna *et al*.,^11^ who reported preferential binding of diruthenium compounds to these same sequence regions. This quantitative agreement between simulations and experiments provide strong support for identifying the cationic form as the biologically relevant binding species.

**FIG. 4.**
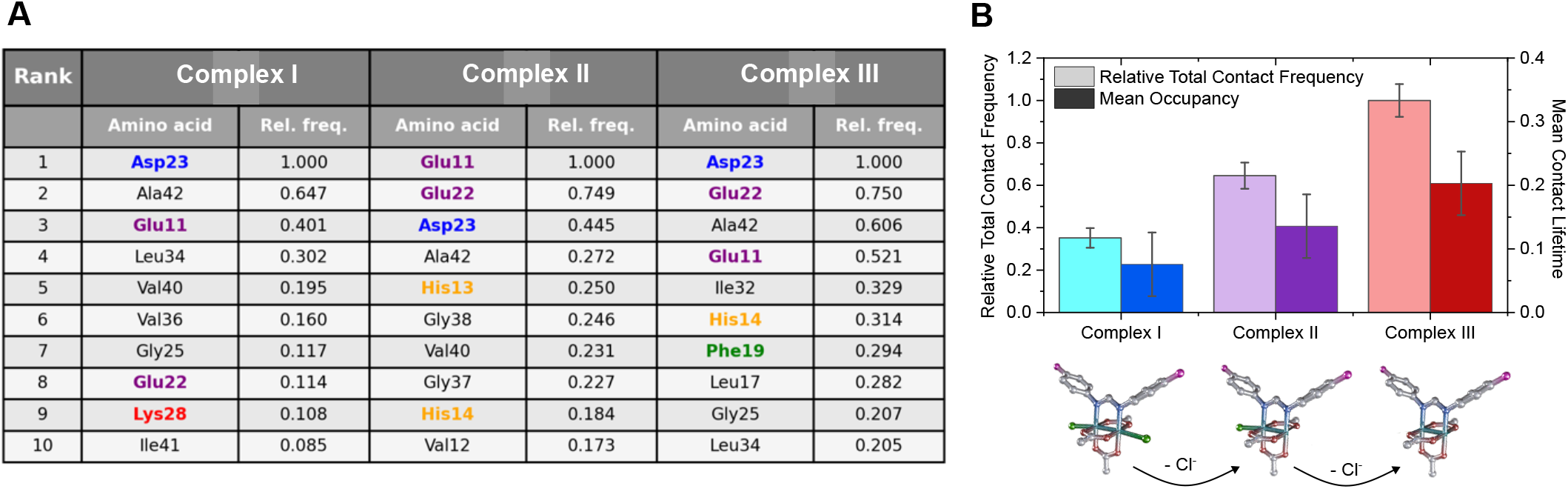
Quantitative analysis of A*β*_42_ interactions with diruthenium complexes. (A) Ranking of the top ten A*β*_42_ residues showing the highest relative contact frequency with Complex I, Complex II, and Complex III. Relative frequencies are normalized by the most interacting residue in each case. (B) Comparison of the relative total contact frequency (light bars, left axis) and mean contact lifetime (dark bars, right axis) for each diruthenium complex. Error bars represent statistical uncertainty across simulations. Structural representations of the three complexes are shown below, highlighting the progressive chloride dissociation from Complex I to Complex III.

Beyond the spatial distribution of contacts, Figure 4B reveals a systematic increase in both the relative total contact frequency and the mean contact lifetime across the series Complex I *→* Complex II *→* Complex III. Complex III not only forms the greatest number of contacts with the fibril surface, but also maintains them for significantly longer duration compared to the neutral or anionic species. This increased contact persistence reflects the formation of a more stable and kinetically robust binding mode, consistent with the broader and more chemically diverse engagement described above. The progressive chloride dissociation illustrated in the structural representations below Figure 4B can thus be interpreted as a charge-activation mechanism: each successive loss of an axial chloride ligand increases the electrostatic affinity of the complex for the fibril surface, expands its binding footprint, and prolongs the lifetime of the resulting complex–fibril encounter. Collectively, these results establish Complex III as the most effective binder within the series, combining broad residue-level engagement with the thermodynamically and kinetically most stable interaction mode.

### Mechanistic basis of fibril destabilization by Complex III: covalent engagement of Asp23 and thermodynamic consequences

Having established that Complex III displays the broadest binding footprint and the most persistent contacts with the A*β*_42_ fibril surface, we investigated the mechanistic consequences of its interaction with Asp23— the residue that consistently ranks as the highest-frequency contact partner across the charge series.

The proposed reaction mechanism, illustrated in Figure 5A, proceeds through sequential ligand substitution steps. In aqueous solution, both axial positions of Complex III are expected to be occupied by water ligands, as reported for related species.^18^ Upon approach of Complex III to the fibril surface, the carboxylate group of Asp23 displaces a coordinated water molecule from an axially labile position of the Ru_2_ core; and direct substitution of that water ligand by the Asp23 carboxylate oxygen then yields a covalently bound adduct in which the fibril residue becomes incorporated into the metal coordination sphere. The lability of axial positions in paddlewheel diruthenium complexes toward displacement by oxygen- and nitrogen-donor biomolecules is firmly established.^14,51^ Moreover, a bond between an Asp residue and an axial position of the Ru_2_ core was observed in the crystal structures obtained from the reaction of [Ru_2_Cl(DPhF)(O_2_CCH_3_)_3_] and [Ru_2_Cl(DPhF)_2_(O_2_CCH_3_)_2_] (DPhF^*−*^ = *N,N’* - diphenylformamidinate) with HEWL.^18^ The recurrence of Asp carboxylates as equatorial or axial anchors across multiple complexes and two structurally distinct model proteins supports the mechanistic hypothesis that Asp23 of the A*β*_42_ fibril occupies the same coordination role here, with the cross-*β* surface architecture presenting this residue in a solvent-exposed, geometrically accessible configuration that favors encounters with the Ru_2_ core.

**FIG. 5.**
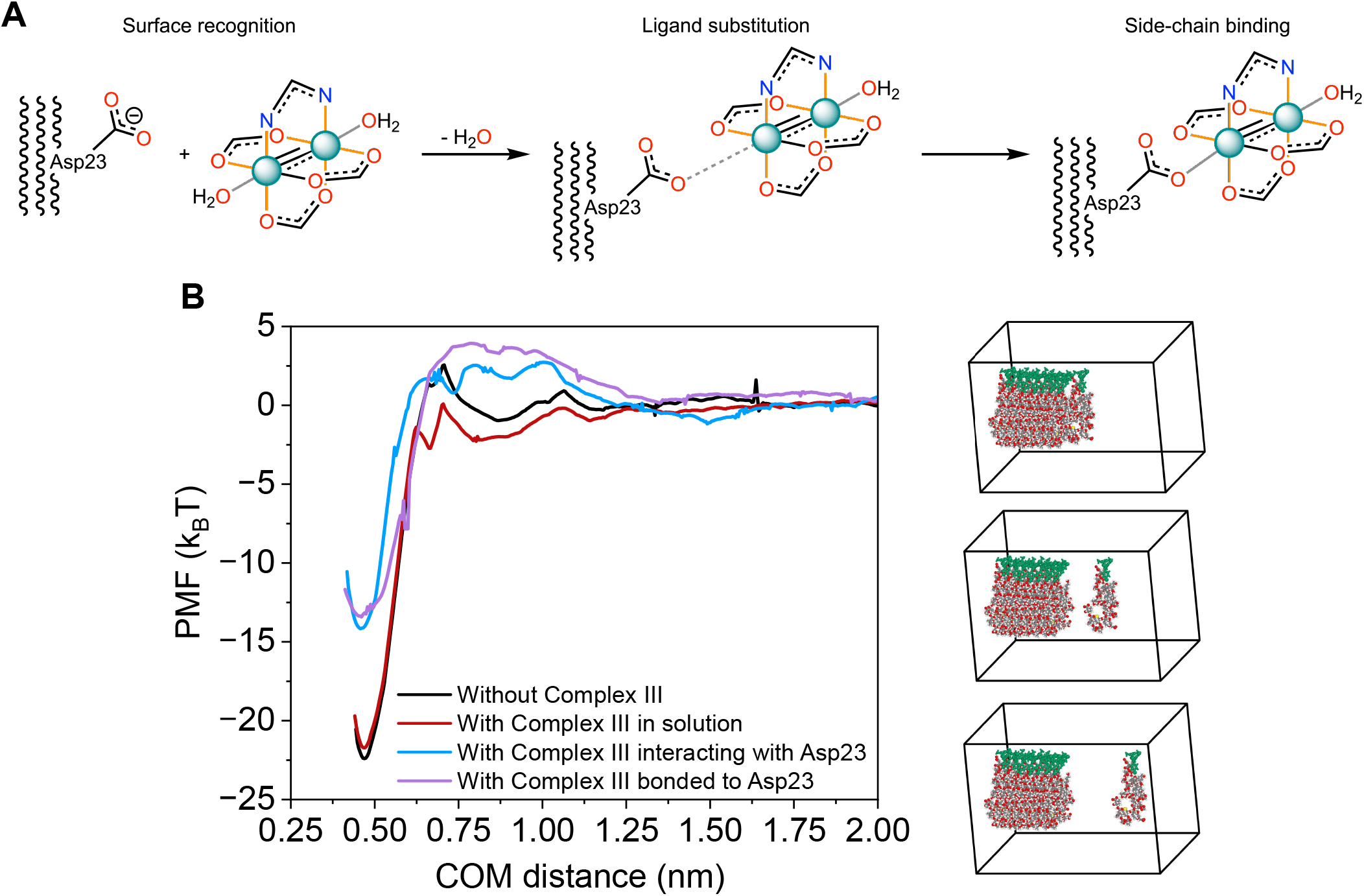
(A) Schematic representation of the proposed mechanism between Complex III and residue Asp23. (B) Potential of mean force (PMF) as a function of the center-of-mass (COM) distance for the different systems: without Complex III (black), with Complex III in solution (red), with Complex III interacting with Asp23 (blue), and with Complex III covalently bonded to Asp23 (purple). (C) Representative simulation snapshots corresponding to the bonded Complex III–Asp23 system (purple curve in panel B; Complex III depicted in green), illustrating the reaction coordinate employed in the PMF calculations and the progressive separation of the fiber along the pulling direction.

To quantify the thermodynamic consequences of this progressive engagement, we computed PMF profiles as a function of the COM distance between one terminal chain and the remainder of the fibril under four distinct conditions: (i) in the absence of Complex III, (ii) with Complex III present in solution but not interacting with the fibril, (iii) with Complex III electrostatically interacting with Asp23, and (iv) with each Complex III covalently bonded to each Asp23 (Figure 5B).

The presence of Complex III in solution without direct fibril contact (Figure 5B, red line) produces only a modest perturbation of the PMF profile relative to the unperturbed system (i.e., in absence of Complex III; black curve), confirming that non-specific solvation effects are limited. In this case, the free-energy minimum remains comparable to that of the native fibril, indicating that the overall thermodynamic stability of the assembled state is largely preserved. However, at intermediate separation distances, the free-energy barrier associated with resolvation of the dissociating chain becomes reduced in the presence of the complex, suggesting that the ligand facilitates solvent penetration and partially destabilizes interchain packing across the reaction coordinate, thereby making fibril dissociation more favorable.

When Complex III interacts electrostatically with Asp23 (Figure 5B, blue line), the PMF profile changes substantially relative to both the native fibril and the “in-solution” condition. The free-energy minimum becomes significantly shallower, indicating a marked reduction in interchain cohesion upon direct interaction of the complex with the fibril surface. This result suggests that non-covalent association with Asp23 driven by electrostatic interactions is already sufficient to destabilize the fibrillar assembly by perturbing the local intermolecular packing and electrostatic balance. In the case of covalent bonding of Complex III to Asp23 (Figure 5B, violet line), the same trend is further amplified. The PMF well becomes even shallower than in the native case, reaching the lowest depth among all systems studied, but similar to the case when Complex III interacts electrostatically with Asp23. The representative simulation snapshots shown in Figure 5C illustrate the progressive structural disruption associated with the covalently bonded system along the pulling coordinate. The sequential images reveal that chain removal in the presence of the covalently bound complex is accompanied by a more pronounced deformation of the fibril terminus, consistent with a reduced cooperative barrier to dissociation.

Taken together, these results delineate a two-stage destabilization mechanism: Complex III first engages the fibril surface through charge-driven, multi-site non-covalent contacts that perturb the local packing of key structural residues, and subsequently forms a covalent adduct with Asp23 that further stabilizes a disruptive interfacial arrangement, inducing reduced thermodynamic stability of the cross-*β* architecture. This dual noncovalent/covalent mode of action represents a mechanistically distinctive feature of the diruthenium scaffold and provides a rational basis for its experimentally observed inhibitory activity toward A*β*_42_ fibril formation.

## CONCLUSIONS

In this work, we have used molecular dynamics simulations to characterize the interaction of a series of paddlewheel diruthenium complexes with A*β*_42_ fibrils at all-atom level resolution. An intermolecular contact map analysis of the A*β*_42_ fibril structure reveals that its stability is governed by a cooperative network of hydrophobic and aromatic interactions, with the C-terminal segment Ile31–Met35 constituting the principal structural core and the central hydrophobic cluster around Leu17– Phe19 providing essential complementary stability (Figure 2). The PMF profile suggests that single-chain dissociation from the terminus of a small fibril entails a free energy cost of approximately 22 *k*_B_*T* (Figure 2), consistent with the high thermodynamic stability of amyloid assemblies^23,49,53^.

Systematic charge variation in diruthenium Complexes I–III from -1 to +1 reveals that electrostatic complementarity is the primary determinant of the binding footprint with the A*β*_42_ fibril surface. As the charge becomes positive, the interaction pattern evolves from a localized engagement around Asp23 to a broad, persistent contact network involving charged, aromatic, and hydrophobic residues across the entire A*β*_42_ sequence, in close agreement with experimentally determined binding sites.^11^ This agreement supports that cationic complexes with axial-free or axial-labile positions are the most biologically relevant fibril-binding species.

Finally, PMF calculations under progressive binding conditions suggest a multistage engagement mechanism rather than a sharp transition between non-covalent and covalent states. Initial electrostatic engagement of Complex III with Asp23 promotes stable association with the fibril surface and locally perturbs intermolecular contacts within the aggregate. Subsequent coordination of the Asp23 carboxylate to the diruthenium core preserves a similar free-energy minimum, with no additional metastable minima emerging along the dissociation coordinate, suggesting that covalent attachment does not substantially strengthen binding affinity itself, but instead stabilizes the complex in a specific interfacial configuration. The principal mechanistic effect therefore appears to arise from the persistent occupation and local reorganization induced by the metal center at the Asp23 region, rather than from a dramatic energetic gain upon covalent bond formation. Such complex-Asp23 engagement enhances A*β*_42_ solubility and lowers the free energy penalty for fibril dissociation.

Together, these findings identify axial chloride dissociation as a charge-activation process that enhances the fibril binding, while covalent coordination acts mainly as a structural locking step that directly interferes with the cooperative interactions sustaining the cross-*β* architecture. The diruthenium scaffold thus combines broad electrostatic surface recognition with site-specific coordination chemistry, providing a mechanistically interpretable mode of action with A*β*_42_ fibrils. More generally, this work establishes a computational framework that links charge state, ligand exchange dynamics, and free-energy profiles as key determinants of metal–fibril interactions, providing a systematic basis for rationalizing and predicting the behavior of charge-tunable metal complexes in amyloid systems.

## Data availability

We provide the relevant data in the repository (GitHub link for the repository) to facilitate reproducibility of our results.

## ACKNOWLEDGMENTS

A. F. acknowledges funding from the Ramon y Cajal fellowship (RYC2021-030937-I) and Spanish National Grant (PID2022-136919NA-C33). A. R. T. acknowledges funding from the European Union Horizon 2020 research and innovation programme (grant agreement 803326 to R.C.-G.). J. R. E. acknowledges funding from Emmanuel College, the University of Cambridge, the Ramon y Cajal fellowship (RYC2021-030937-I), the Spanish scientific plan and committee for research reference PID2022-136919NA-C33, and the European Research Council (ERC) under the European Union’s Horizon Europe research and innovation program (grant agreement no. 101160499). S. H. acknowledges the Regional Ministry of Education, Science and Universities of the Community of Madrid (TEC-2024/TEC-85). A. T. acknowledges the Juan de la Cierva fellowship grant (JDC2024-053530-I) funded by the Spanish Ministry of Science, Innovation and Universities (MICIU) and the State Research Agency (AEI). A.M. thanks MIUR PRIN 2022 - Cod. 2022JMFC3X, “Protein Metalation by Anticancer Metal-based Drugs” for financial support. A.O. acknowledge funding from CRIS Cancer Foundation (AOF.C01CRIS and AOF.M01CRIS).

